# Borrowing ecological theory to infer interactions between sensitive and resistant breast cancer cell populations

**DOI:** 10.1101/2022.02.18.481041

**Authors:** Zachary Susswein, Surojeet Sengupta, Robert Clarke, Shweta Bansal

**Affiliations:** Department of Biology, Georgetown University, Washington, DC, USA; The Hormel Institute, University of Minnesota, Austin, MN, USA

## Abstract

While some forms of breast cancer are highly responsive to treatment, endocrine therapy-resistant breast cancers are disproportionately lethal. There has been significant progress in understanding how endocrine therapy-resistant strains evolve from therapy-susceptible strains of cancer, but little is understood about the proliferation of resistance through cancer cell populations, or the interactions that occur between populations of resistant and sensitive cells. In this study, we characterize the nature of the ecological interaction between populations of resistant and susceptible breast cancer cells to reveal novel methods of controlling drug resistance.

Using in-vitro data on fluorescent-tagged resistant and susceptible cells, we use an image processing algorithm to identify and count cell growth till equilibrium. We then borrow theory from population biology to infer the type of ecological interaction that occurs between populations of resistant and sensitive cells. In particular, we use a Bayesian approach to fit single culture cell populations to infer density-dependent growth parameters (growth rate, carrying capacity) and a Generalized Lotka-Volterra model to understand how susceptible and resistant co-culture populations may be depressing or supporting growth of the other.

Our results identify a net mutualistic interaction between the susceptible and resistant cancer strains, demonstrating that there are ecological dynamics to cancer resistance. Our findings also suggest that ecological dynamics change in the presence of therapy, and that an adaptive treatment protocol can induce cycling behavior suggesting that heterogeneous ecological effects contribute to empirically observed adaptive-therapeutic dynamics.

## Introduction

Breast tumors that express the estrogen receptor alpha (ESR1; ER) often respond to an initial endocrine therapy in the form of either an aromatase inhibitor or an antiestrogen. Despite the effectiveness of these therapies, proliferation of acquired or de novo resistant cells can diminish tumor response to treatment, increasing mortality and often necessitating the use of cytotoxic therapies, especially in disseminated cancers [1, 2]. Traditional treatment approaches can speed the emergence of therapy resistance through evolutionary pressure, selecting for therapy-resistant cells by eliminating those that are therapy-susceptible [3, 4, 5]. Based on the “maximum tolerated dose” principle, this strategy focuses on diminishing tumor burden by eliminating as many susceptible cells as possible. However, when recurrence occurs, less effective or poorly tolerated systemic therapies are used to eliminate the cancer cells that are now resistant to the first-line therapy [1, 3, 6]. Nonetheless, recurrent breast cancer remains largely incurable. In contrast, the adaptive therapy paradigm focuses on limiting future tumor growth or proliferation, aiming to decrease time to progression without necessarily reducing tumor size [3]. This approach takes advantage of ecological and evolutionary dynamics, including the spatial heterogeneity of resistant phenotypes across the tumor and the fitness cost of resistance to find the optimal therapy to keep tumor growth in check for the longest possible time [4, 7, 8, 9, 10]. By using strategies like dose titration and “dose skipping” such as treatment vacations or drug cycling, adaptive therapy approaches can take advantage of intratumoral competition to limit overall tumor growth and spread of the resistant phenotype [11, 7, 12, 3, 13]. Emerging research further refines these approaches, applying evolutionary principles to optimize individual-specific dose titration across time to minimize the development and spread of therapy resistance [12, 14, 15].

Mathematical models are used in the adaptive therapy literature to explore ecological and evolutionary dynamics within tumors; these models frequently rely on two key assumptions: the fitness cost of resistance and robust competition between intratumor cell populations [3, 15]. Assuming a fitness cost to resistance, a resistant strain is only more fit than a susceptible strain in the presence of therapy. Additionally, under competition between strains, a more fit strain will reproduce more quickly in the environment in which it is more fit, outcompeting other strains. By taking advantage of this ecological dynamic, adaptive therapy aims both to prevent the spread of the resistance and to limit the evolution of new resistant genotypes. A range of complex modeling strategies build on this approach, using mechanistic models of tumor dynamics to examine the evolution of the resistant phenotype, to develop strategies that minimize resistance proliferation, and to optimize treatment approaches that maximize competitive dynamics within the tumor [16, 17, 18, 19, 20].

Despite the ubiquity of these assumptions, a full understanding of the interpopulation ecological dynamics between cancer phenotypes, including the magnitude of competition and the potential fitness cost of resistance, has not yet been developed [21, 22, 10]. Indeed, cooperative interactions between sub-populations of heterogeneous tumors are well-described, both *in vitro* and *in vivo*, calling into question the validity of the intratumor competition assumption [23, 24, 22, 25]. Mathematical models have been used to identify not only mechanisms driving competition for scarce resources (see e.g. [26, 17, 7, 27]), but also pro-growth (mutualistic) interactions between intratumor strains – fostering greater overall tumor expansion [22, 28]. Similarly, resistance mechanisms can maintain or even increase fitness in the absence of therapy, challenging the fitness cost of resistance assumption [29].

Here, we examine the fitness cost of resistance and the interaction between sensitive and resistant populations, fitting classical ecological models to *in vitro* cell culture population data. Our Bayesian mechanistic approach allows for flexible model specification, rich parameterization of uncertainty, and direct interpretation of model coefficients. Using longitudinal observations of cell abundance in the presence and absence of therapy, we quantify growth parameters and net interactions between the populations. Our results suggest that ecological dynamics change in the presence of therapy and that assumptions of a fitness cost to resistance and inter-population competition should be verified empirically on a system-by-system basis. We also apply an adaptive treatment protocol to our mechanistic model, finding that it can decrease the resistant population size and induce population cycling behavior – despite the absence of a fitness cost to resistance – suggesting that heterogeneous ecological effects contribute to empirically observed adaptive-therapeutic dynamics.

## Results

Here, we examine the type and magnitude of the ecological interaction between therapy-sensitive (LCC1) and therapy-resistant (LCC9) populations. We culture LCC1 and LCC9 populations in monoculture and in coculture, count the cell population size every two hours, and fit logistic growth and Lotka-Volterra growth models, respectively, to the temporal growth dynamics. We execute this procedure for both the control and treated conditions, estimating growth parameters and inter-population interaction strength in each case. We simulate from these posterior distributions to examine the impact of inter-population interactions on predicted growth.

### In-vitro cell populations can be rigorously quantified

We develop an image processing algorithm to systematically quantify cell population sizes over time. Due to the noisy nature of the cell culture experiments, traditional automated tools, such as ImageJ, are unable to differentiate between cells and background noise in this system. These images present a range of issues for rigorous quantification, including fluctuations in color density across the camera field, bunching in cell density (i.e. many groups of cells tightly clumped together), and changes in brightness over time. Using the scikit-image library, we develop a robust image processing pipeline that rapidly and accurately quantifies cell abundance. We validate these algorithmic counts against manual image counts, finding high agreement between the two (Pearson’s *ρ* = 0.97; Figure 1). Accurate quantification of abundance enables systematic characterization of population dynamics over time.

**Figure 1:**
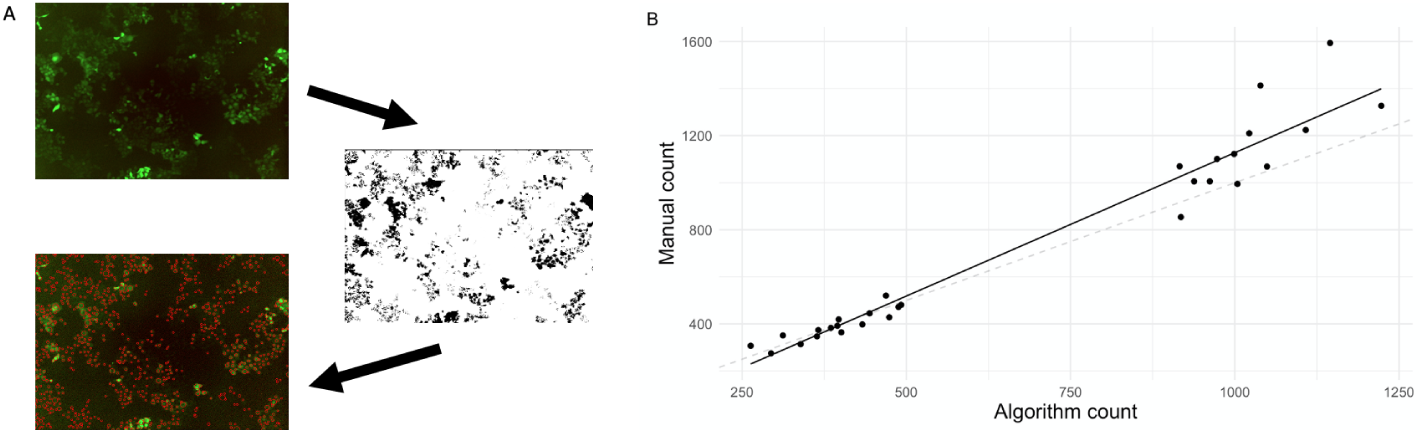
Automated counting algorithm produces counts of in vitro cell population sizes. A) Our automated algorithm uses a two-stage process to reduce image noise through a sigmoid correction, adaptive thresholding, artifact removal, and cell identification. Contrast and intensity corrections reduce background noise and normalize cell intensity, allowing for more accurate cell identification. B) Comparison of algorithm counts to manual counts with simple linear regression (solid black line) indicates satisfactory cell counts from images. Pearson’s correlation coefficient of 0.97 (95% CI: 0.94, 0.99) shows that algorithm and manual counts are highly correlated. Higher cell counts are estimated as slightly larger by the algorithm count than by manual count (dashed line one-to-one).

### Resistance does not confer a fitness cost

Before characterizing interactions between populations, we independently characterize the parameters governing growth dynamics of therapy-sensitive and therapy-resistant populations. We employ the logistic growth model, fitting population dynamics parameters to the data quantified through our cell counting algorithm. For each condition, sensitive or resistant cell line and in the presence of vehicle or treatment, we fit a separate logistic growth curve, finding a condition- and strain-specific intrinsic growth rate (*r*) and carrying capacity (*K*) (Figure 2A). Posterior estimates for sensitive (LCC1) growth in the presence of treatment are lower than those in the presence of vehicle, with estimated *r* and *K* both decreased – consistent with the expected effects of treatment on the sensitive phenotype (Figure 2B). Posterior estimates of resistant (LCC9) growth parameters, both with vehicle and with treatment are similar to, but slightly higher than, those from the sensitive vehicle condition. In this system, the resistant population is expected to grow at least as quickly as the susceptible population in monoculture regardless of treatment regime; there is not a fitness cost of resistance.

**Figure 2:**
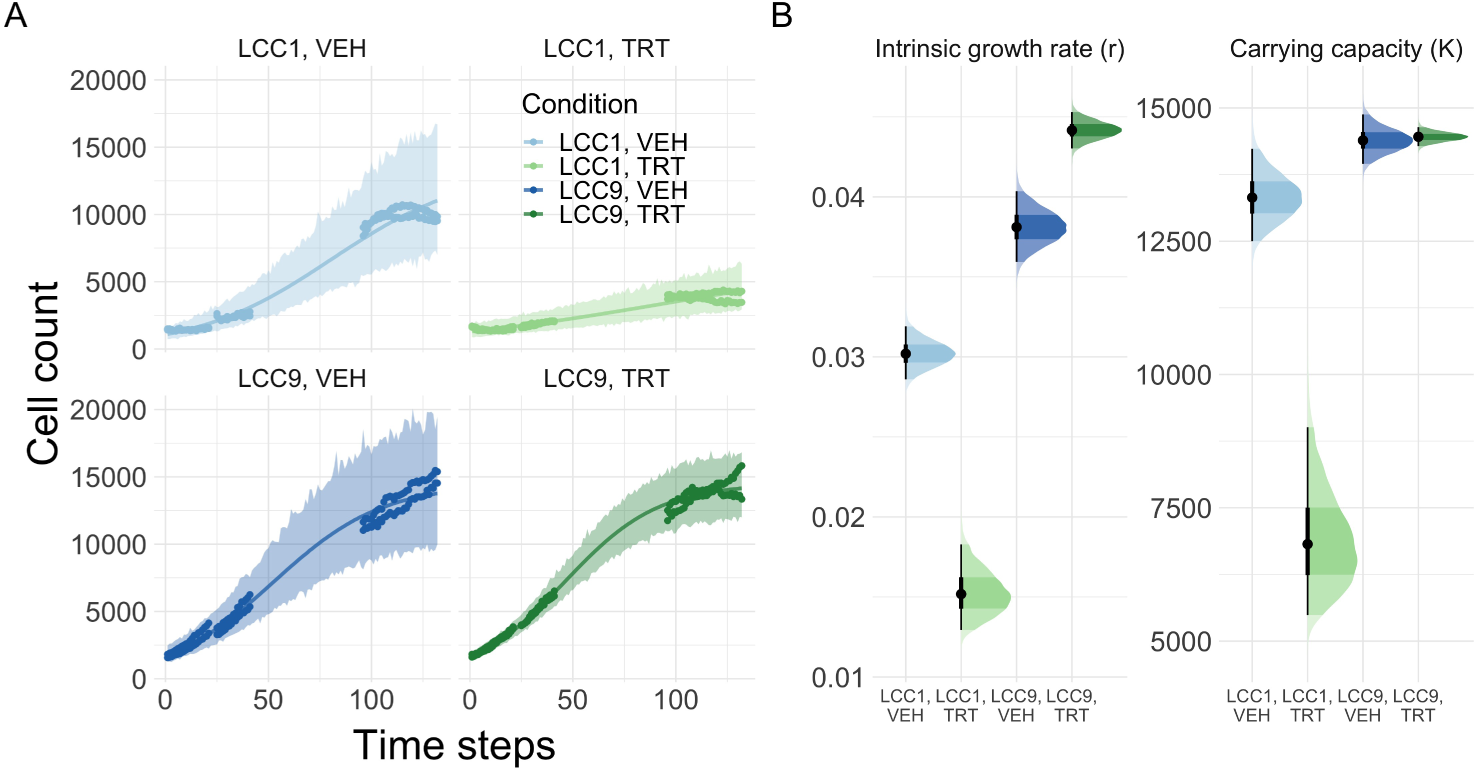
Resistant (LCC9) population demonstrates no fitness cost of resistance compared to susceptible (LCC1) population. A) Bayesian posterior predictive distributions of logistic growth model to *in vitro* monocultures show heterogeneous growth dynamics. Logistic growth model posterior mean (solid line) and 95% curve-wise credible interval (shaded interval) reproduce observed cell cultures (points) across time. B) Treatment diminishes sensitive (LCC1) cell population growth in monoculture, while the resistant (LCC9) cell population is unaffected. Sensitive intrinsic growth rates (*r*) and carrying capacities (*K*) are decreased by therapy, while resistant intrinsic growth rates do not change substantially and are at least as high as those of sensitive populations in all cases, as measured by the 50% and 95% credible interval. Note that VEH corresponds to vehicle and TRT corresponds to treatment.

We use the Lotka-Volterra competition model to quantify cell growth in co-culture, using the monoculture posterior distributions as informative prior distributions for growth parameters. Sensitive and resistant populations in the co-culture vehicle condition do not grow more slowly than their respective mono-culture counterparts, which suggests that intercellular signaling dynamics are supporting the increased cellular abundance. Indeed, posterior estimates of intrinsic growth rates and carrying capacities are unchanged from monoculture estimates; therefore, any difference in growth is driven by ecological interaction between sensitive and resistant populations in co-culture, not changes in condition-specific growth parameters (Figure 3B). For the populations in co-culture under treatment (both LCC1 and LCC9) each grows more slowly than it does alone, in the monoculture condition, or together, in the co-culture vehicle condition (Figure 3A). However, the growth parameter estimates are again unchanged from those in monoculture. Therefore, any heterogeneity in growth dynamics across the treatment conditions is driven by heterogeneity in ecological interaction between the populations.

**Figure 3:**
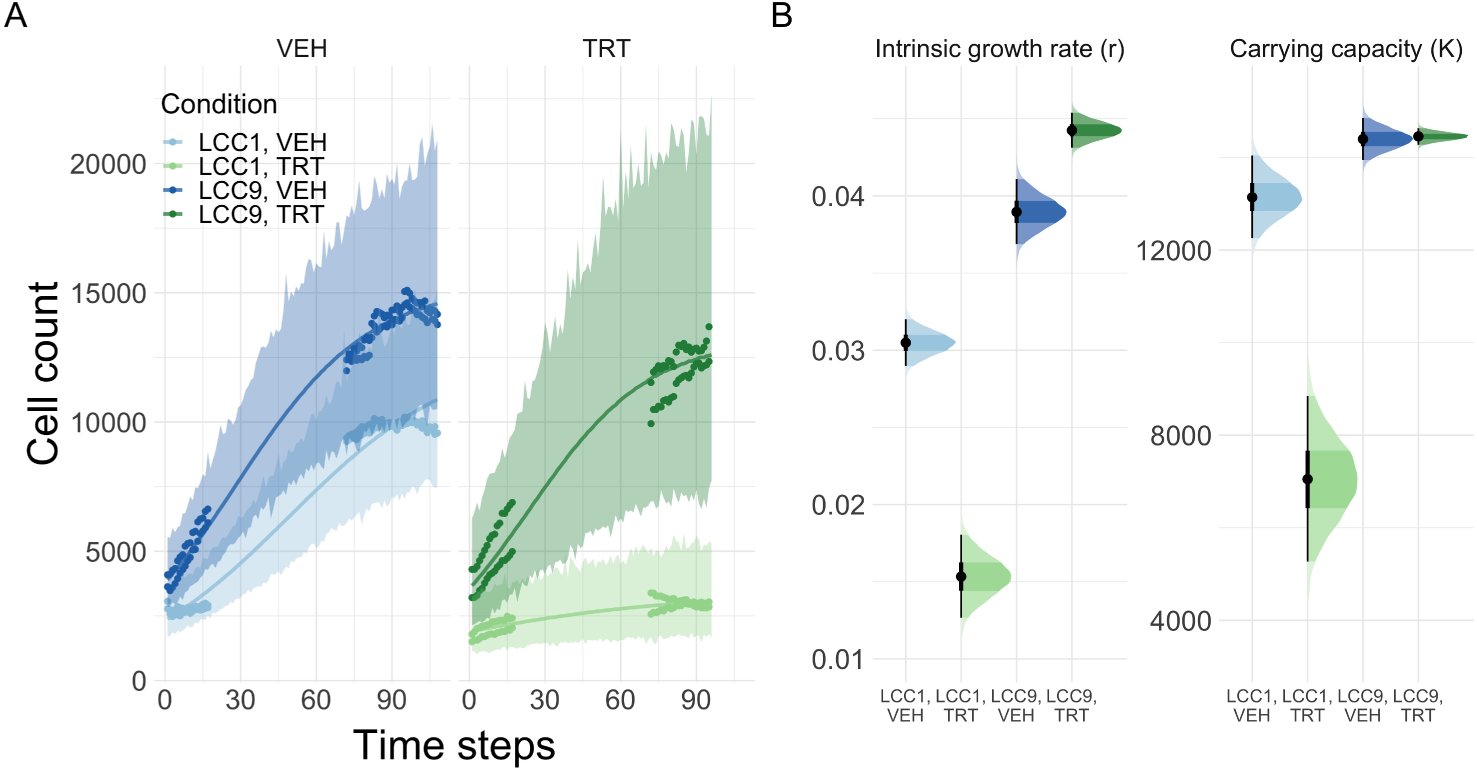
Posterior estimates of sensitive (LCC1) and resistant (LCC9) populations grown in coculture in two replicates are similar to those from monoculture. A) Bayesian posterior predictive distribution of the Lotka-Volterra competition model to coculture of sensitive and resistant cell populations shows accurate recovery of growth dynamics. Lotka-Volterra model posterior mean (solid line) and 95% posterior curve-wise credible interval (shaded interval) reproduce observed cell cultures (points) across time. B) Posterior 50% and 95% credible intervals for intrinsic growth rates and carrying capacities reproduce logistic growth estimates. Because growth parameters are unchanged from the mono culture posterior estimates, accurate coculture fits are driven not by changes in growth parameters, but by the net ecological interactions. Note that VEH corresponds to vehicle and TRT corresponds to treatment.

We identify heterogeneous ecological interactions between therapy-sensitive and therapy-resistant strains across treatment conditions, suggesting that treatment alters cross-strain signaling interactions. Ecological interactions between sensitive and resistant populations grown in co-culture are only weakly competitive and heterogeneous across each treatment condition (Figure 4). If a strong and symmetric competition existed between sensitive and resistant populations, the net ecological interaction coefficients (*α* and *β*) would be at least equal to 1, with the inter-population reduction in available carrying capacity reducing by one per-individual – equivalent to the assumed reduction caused by intra-population competition [30]. Instead, *α* and *β* are substantially below 1 in both the vehicle and treatment conditions. Thus, the net ecological interactions are overall less competitive because of pro-growth interactions between the populations (Figure 4). The net ecological interactions captured by *α* and *β* are near 0 in the presence of vehicle – co-culture growth of the two populations is only weakly altered by inter-population interaction. While the magnitude of these interactions approaches zero, their sign can fluctuate across sensitivity analyses and should not be too closely interpreted (INSERT SUPP FIG). These interactions are heterogeneous across treatment conditions, with *α* and *β* between 0 and 1; each additional cell in one population reduces the available carrying capacity of the other population by between 0.1 and 0.7 cells (based on their respective credible intervals). While more competitive than with vehicle treatment alone, the net ecological interactions with treatment again place almost all posterior density for *α* and *β* below 1, with pro-growth interactions between the populations reducing overall competition.

**Figure 4:**
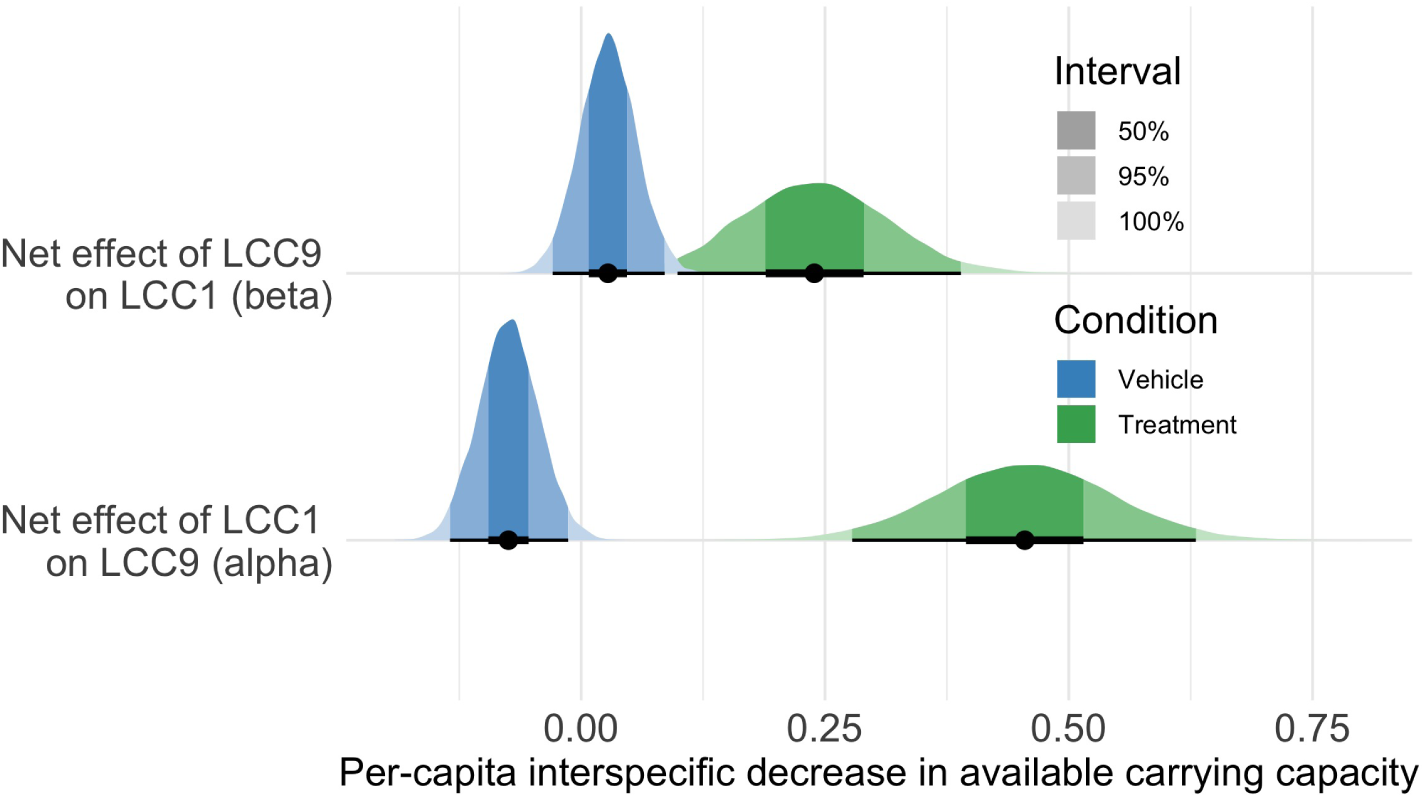
Net ecological interactions between sensitive (LCC1) and resistant (LCC9) populations are only weakly competitive and heterogeneous across treatment conditions. Ecological interaction coefficients (*α* and *β*) measure the overall effect of all interactions between the sensitive and resistant populations, including pro-growth signaling and competition for scarce resources. In the absence of treatment (i.e., vehicle), mutualism and competition are closely balanced, with growth remaining at near-monoculture levels (*α* ≈ *β* ≈). With treatment, net interactions are more competitive; each population is expected to reduce the other population’s carrying capacity per cell by between 0.20 and 0.50 cells. The probability of either interaction coefficient equaling 1 is close to zero, suggesting that inter-population competition is weaker than intra-population competition.

### Heterogeneous ecological effects alter population dynamics

The observed heterogeneity ecological dynamics has important implications for coculture growth dynamics, altering predicted growth patterns under varying treatment regimes. A metronomic therapy regimen alternating between vehicle and treatment dynamics induces cycling behavior in the system, with sensitive and resistant populations both decreasing in the presence of treatment because of the heterogeneous ecological dynamics across populations and treatment conditions (Figure 5). Importantly, this dynamic is driven not just by changes in relative amounts of inter-population competition, but also by changes in population-specific fitness under the different treatment conditions (i.e. *r, K*). Despite the absence of a fitness cost of resistance in this system, both populations persist and neither is outcompeted. However, assuming robust competition between the populations (setting *α* = *β* = 1) alters dynamics such that the sensitive population is rapidly outcompeted (Figure 5B). Assuming stronger competition than is actually occurring substantially alters predicted dynamics.

**Figure 5:**
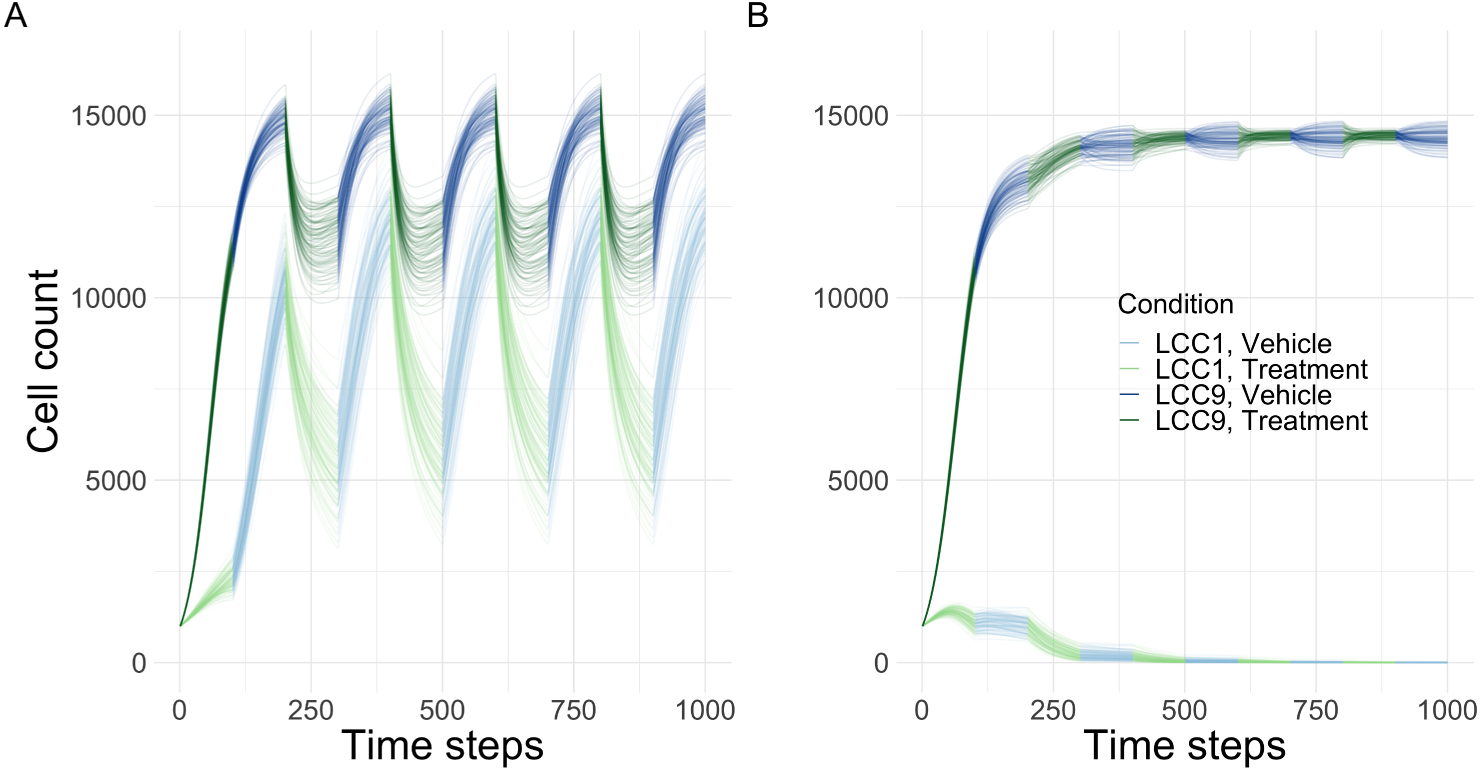
Ecological dynamics alter predicted coculture growth. A) Predicted growth *in silico* using 100 realizations from the posterior distributions of the Lotka-Volterra coculture models with 5 treatment cycles – alternating between vehicle and treatment growth models. Despite the lack of a fitness cost of resistance, changes in LCC1 carrying capacity and ecological interactions between populations induce cycling behavior in the system. B) Predicted growth in silico from 100 posterior realizations and net ecological interaction fixed at robust symmetric competition between populations, *α* = *β* = 1. By assuming stronger competition than actually occurring in this system, sensitive populations are predicted to be outcompeted and rapidly decline despite the alternating treatment conditions.

## Discussion

Tumors are complex ecological environments, composed of heterogeneous interacting populations. In this work, we show that assumptions of a fitness cost of resistance and strong inter-population competition between therapy-sensitive and -resistant populations can be oversimplifications and lead to inaccurate predicted dynamics. As we show, the ecological dynamics between populations can vary with treatment conditions, altering growth dynamics. This change is driven by the interaction between the populations, not a change in the growth parameters (relative to monoculture). We do not attempt to generalize our quantitative results outside of our specific *in vitro* system (see e.g. [31]), but argue that heterogeneity in ecological dynamics across treatment conditions can contribute to the identified population cycling behavior in empirical adaptive therapeutic systems (see e.g. [17, 14]) and that pro-growth interactions can suppress competitive dynamics more than commonly assumed in the adaptive therapy literature.

Mathematical modeling of adaptive therapy often includes the impact of inter-population competition, either through the arrest of growth when resource constrained (spatially constrained) in lattice-based or agent-based frameworks (e.g. [7, 28]) or through the use of a mean-field approximation averaging over the impact on the population as a whole (e.g. [32, 17, 29]). The Lotka-Volterra model framework is a popular implementation of the latter approach, providing interpretable coefficients and a strong basis in ecological theory [30]. In Zhang et al., the Lotka-Volterra model is used for three populations, with a fitness cost of resistance assumed and competition assumed to be stronger than we identify here, showing that adaptive therapy can induce oscillations in tumor burden that delay time to progression [32]. Strobl et al. relax the fitness cost of resistance assumption but assume robust competition (set *α* = *β* = 1), finding that adaptive therapy can limit time to progression even without a fitness cost of resistance [29]. In our system, we experimentally identify stronger pro-growth mutualistic interaction between sensitive and resistant populations than commonly assumed, but show that these can still contribute to cycling dynamics – even without a fitness cost of resistance. Furthermore, we carefully quantify the uncertainty in this empirical system, fitting full posterior distributions to the observed data. With this uncertainty, we quantify the range of predicted outcomes under different treatment regimens – not just the expected mean behavior.

In the posterior predictions from the fitted differential equation model, the predicted cycling behavior is caused not just by the ecological interaction between populations, but also by the relative fitness of the susceptible population in the absence of therapy. The change in susceptible carrying capacity between the treatment and vehicle conditions allows the susceptible population to rapidly increase during treatment cycling, with resistant populations decreasing to compensate because of the weakly competitive interaction between the populations. While the carrying capacity is commonly nondimensionalized when working with the Lotka-Volterra equations to ease computation, doing so can potentially decrease biology accuracy when the true carrying capacity is unknown or differs across conditions. These results suggest that therapeutic treatment models that do not account for the sources of variability in population response to treatment may improperly estimate the impact of therapies or the relative contribution of potential mechanisms.

Understanding precisely how the interactions between sensitive and resistant cells occur, and how these are affected by treatment, are beyond the scope of the present study. Nonetheless, individual cancer cells in a population have several well-established means to communicate. The most widely studied mechanisms involve the secretion of materials (paracrine communication) as free molecules such as secreted growth factors or cytokines [33, 34, 35]. Growth factors in the tumor microenvironment can induce ligand-independent activation of ER [36, 37], conferring endocrine resistance [33, 34]. Paracrine communication also includes the packing of intracellular materials into microvesicles that are then released into the microenvironment and beyond [38, 39, 40]. The cargo of microvesicles can be complex and includes small molecules such as amino acids, simple sugars, and non-coding RNAs (ncRNAs) but also larger macromolecules such as lipids, mRNAs and cDNAs [41]. The ability to communicate regulatory molecules such as ncRNAs or cDNAs that could be incorporated into the genome, could allow for the target cells to either temporarily or permanently acquire features of the cell from which the microvesicles originated.

Since cells are often packed tightly together in the tumor microenvironment, this cell-cell contact also enables juxtacrine communication. Juxtacine communication can arise from adjacent protein-protein interactions, such as where the extracellular membrane of one cell expresses a receptor that binds a ligand in the extracellular membrane of it neighbor. Cells can also directly share components of their respective cytosols through gap junctional intercellular communication facilitated by pores formed by the connexins. GJIC generally allow for the sharing or relatively small molecules such as amino acids, glucose, and second messengers like Ca++, IP3, and cAMP [42, 43, 44, 45].

*In vitro* systems are idealized, ignoring many complexities of *in vivo* tumor dynamics, but the robust literature on intratumor mutualism suggests that mutualistic dynamics persist outside of idealized settings [22]. Our results suggest that this mutualism can be strong enough to completely counter inter-population competition in idealized conditions, suggesting that mutualism cannot be ignored by adaptive-therapeutic modeling. As we show, systems lacking both a fitness cost to resistance and robust competition are able to produce oscillations in resistant population size – suggesting that these dynamics can contribute to empirical observations. When modeling these systems, assuming robust competition or, indeed, competition at all (i.e. *α, β >* 0) can be a mistake.

The observations here may have clinical relevance. The ER+ breast cancer subtype is characterized by a feature generally referred to as “dormancy”, where many tumors recur a decade or more after what had otherwise appeared to have been a curative intervention with an endocrine therapy. Why these tumors remain dormant and clinically undetectable for so long is poorly understood. Cycling in the growth rate of mixed cell populations could lengthen the time it takes for a population to grow to a clinically detectable size. Similarly, the reduction in carrying capacity could limit the size to which these tumors can grow. Hence, only once a tumor can escape both the growth rate and carrying capacity constraints could it reach a size that would be detected clinically as a breast cancer recurrence. Understanding the molecular features driving these changes in carrying capacity and growth rate remains an area of active research.

## Methods

We grow susceptible (LCC1) and resistant (LCC9) populations in monoculture and in coculture with and without treatment, for a total of eight treatment conditions, and observe growth every two hours. We apply the logistic growth model to cell monocultures, producing estimates of population growth parameters. We apply the Lotka-Volterra model to cell co-cultures, producing estimates of net ecological interactions between populations, in vehicle and in treatment. Using the model-based estimates, we simulate growth in the presence of a metronomic therapy regime.

### Cell culture

LCC1-egfp cells are estrogen-independent and sensitive to 4-hydroxy-tamoxifen (4-OHT) and fulvestrant are derived from LCC1 cells ([46]) that stably express enhanced green fluorescent protein (egfp), while LCC9-mCherry cells are derived from LCC9 cells ([47]) that stably express m-cherry fluorescent protein and are resistant to both 4-OHT and fulvestrant. LCC1-egfp and LCC9-mCherry cells were cultured in phenol red-free, Modified IMEM media supplemented with 5% charcoal-stripped fetal bovine serum. Five thousand LCC1-egfp and LCC9-mCherry cells per well were seeded in a 12-well plate either alone or in co-culture in presence or absence of 500nM fulvestrant. The cells were kept in Incucyte® SX-3 live cell analyzer, at 37°C, housed inside a CO2 incubator. The red and green fluorescent images representing LCC9 and LCC1 cells respectively were captured every two hours during the length of the experiment.

### Image Processing

We process cell culture images from the eight different treatment conditions using scikit-image in Python [48]. We use a two-stage process, using a sigmoid correction and local adaptive thresholding in the initial stage to remove background noise and equalize cell brightness intensity. The second stage locally contrasts and thresholds, removes artifacts, and counts individual cells. There are 8 separate images (one per field) per well per time point; fields are aggregated to the smallest independent unit, the well. There are two replicates for each treatment condition. We validate cell counts on images manually counted by two individuals using simple linear regression.

### Ecological models

To characterize the growth of each population independently, we apply the well-characterized logistic growth model to *in vitro* monoculture growth data. The logistic growth model is expressed as a differential equation:

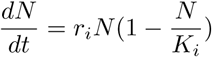

where *N* is the population size at time *t, r*_*i*_ is the condition-specific intrinsic growth rate, and *K*_*i*_ is the condition-specific carrying capacity. The four conditions are LCC1 & vehicle, LCC1 & therapy, LCC9 & vehicle, and LCC9 & therapy. The intrinsic growth rate (*r*) governs a population’s rate of exponential growth in the absence of constraints, inversely proportional to the doubling time; *r* is the sum of the population’s instantaneous birth and death rates without resource constraints [30]. The carrying capacity (*K*) is the maximum population size that can be supported by the environment – it is population- and environment-specific [30].

To characterize the growth of sensitive and resistant populations in co-culture, we use the Lotka-Volterra model, an extension of the logistic growth model for interacting populations. The Lotka-Volterra model extends the logistic growth model with a linear interaction term, allowing for both intra- and inter-population competition [30]. It is represented as a system of paired ordinary differential equations:

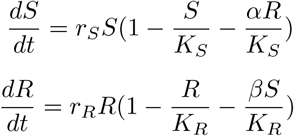

where S and R are the sensitive and resistant population sizes at time *t* respectively, *r*_*i*_ are the intrinsic growth rates, *K*_*i*_ are the carrying capacities, *α* and *β* are the ecological interaction terms, and the impact of intra-population competition is fixed at 1. We fit the model separately for each treatment condition, applying posterior distributions from the logistic growth models of monoculture growth as informative prior distributions on the *r* and *K* parameters in the Lotka-Volterra model. Net ecological interaction terms (*α* and *β*) are assumed by the model to represent a constant mean inter-population effect acting equally on every individual cell. As cells are grown on complete media, the only limiting resource in this *in vitro* system is space, so the competitive mechanism is pre-emptive competition [30].

### Model parameter inference from cell culture data

We fit the ecological models to the cell culture time series with a Bayesian approach using Hamiltonian Monte Carlo. Models are implemented in the R language interface to Stan [49]. We remove the first 48 hours from the time series to remove the period of no growth after initial introduction and remove the last 22 hours from the treatment coculture condition because of a sudden decline in one biological replicate judged to be due to laboratory error. Missing cell culture time points are imputed from the model.

All logistic growth parameters are assigned weakly informative priors to regularize inference. Coculture parameters are assigned informative priors using the posteriors from monoculture growth models except for *α* and *β*, which are assigned standard normal priors. Prior predictive distributions suggest appropriately specified prior distributions [50, 51] We perform HMC sampling with 4 chains with 10000 iterations per chain. In all cases, traceplots indicate the chains are well-mixed, the Gelman-Rubin split 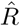 statistic is less than 1.01, and effective sample size is ratio is above 50%, there are no divergent transitions, transitions that exceed maximum treedepth, or transitions with high Bayesian fraction of missing information, suggesting that the chains have converged and there are no numerical issues with model fit. We find that the posterior predictive distributions recover the data, suggesting no systematic errors in model fit. We represent uncertainty in the posterior predictive distributions with 95% curve-wise credible intervals, representing the region with high probability of containing the entirety of the true curve – not just the probability of containing each point along the curve. Cell culture time series, Stan model code, R language analysis code, and original model diagnostics and results are available on Github.

### Treatment simulations

We predict cell population dynamics under a range of treatment conditions. For the treatment cycling condition, we simulate population dynamics under growth and treatment parameterizations for 100 Monte Carlo simulations. In each simulation, we examine 5 treatment cycles, with 100 time points per cycle. We stochastically parameterize growth dynamics by simulating realizations of each growth parameter from normal distributions with the same first two moments as those of the respective marginal posterior distributions. For the condition with specified competition, we fix the ecological interaction coefficients (*α* and *β*) at 1 and repeat the treatment cycling as specified. We solve the system using the ‘ode’ function in the ‘deSolve’ package with the LSODA integrator. We repeat this procedure for 100 Monte Carlo simulations.

## Data availability

Original count data, sample images, image processing code, and analysis code are available at https://github.com/bansallab/breast_cancer.

## Acknowledgements

We acknowledge support from DoD Award W81XWH-18-1-0722,BC171885 for all authors.

